# Small Molecule Inhibitors of the Human Histone Lysine Methyltransferase NSD2 / WHSC1 / MMSET Identified from a Quantitative High-Throughput Screen with Nucleosome Substrate

**DOI:** 10.1101/208439

**Authors:** Nathan P. Coussens, Stephen C. Kales, Mark J. Henderson, Olivia W. Lee, Kurumi Y. Horiuchi, Yuren Wang, Qing Chen, Ekaterina Kuznetsova, Jianghong Wu, Dorian M. Cheff, Ken Chih-Chien Cheng, Paul Shinn, Kyle R. Brimacombe, Min Shen, Anton Simeonov, Haiching Ma, Ajit Jadhav, Matthew D. Hall

## Abstract

The activity of the histone lysine methyltransferase NSD2 is thought to play a driving role in oncogenesis. Both overexpression of NSD2 and point mutations that increase its catalytic activity are associated with a variety of human cancers. While NSD2 is an attractive therapeutic target, no potent, selective and cell-active inhibitors have been reported to date, possibly due to the challenging nature of developing high-throughput assays for NSD2. To establish a platform for the discovery and development of selective NSD2 inhibitors, multiple assays were optimized and implemented. Quantitative high-throughput screening was performed with full-length wild-type NSD2 and a nucleosome substrate against a diverse collection of known bioactives comprising 16,251 compounds. Actives from the primary screen were further interrogated with orthogonal and counter assays, as well as activity assays with the clinically relevant NSD2 mutants E1099K and T1150A. Five confirmed inhibitors were selected for follow-up, which included a radiolabeled validation assay, surface plasmon resonance studies, methyltransferase profiling, and histone methylation in cells. The identification of NSD2 inhibitors that bind the catalytic SET domain and demonstrate activity in cells validates the workflow, providing a template for identifying selective NSD2 inhibitors.

## Introduction

Epigenetic modifiers are widely recognized as targets for therapeutic intervention, due to their critical roles in regulating gene expression and chromatin integrity in addition to their dysregulation in a range of human pathologies. In particular, the NSD (nuclear receptor-binding SET domain) family of histone lysine methyltransferase enzymes, NSD1, NSD2/WHSC1/MMSET, and NSD3/WHSC1L1, have all been implicated as cancer therapeutic targets (1). The catalytic SET domain (Suppressor of variegation, Enhancer of zeste, Trithorax) of NSD family enzymes catalyzes the mono- and di-methylation of the ε-amine of lysine 36 of histone H3 (H3K36), utilizing the methyl donor S-adenosyl-L-methionine (SAM) (Fig. 1A). The non-overlapping roles of the NSD family of enzymes in normal physiology are attributed to a variety of additional domains including a PWWP (proline-tryptophan-tryptophan-proline) domain, PHD (plant homeodomain), zinc finger domains and a high mobility group box (HMG) domain (1).

NSD2 has been implicated as a therapeutic target for a variety of cancers. Because the gene is located within the Wolf–Hirschhorn syndrome (WHS) critical region of chromosome 4, NSD2 is also known as Wolf–Hirschhorn syndrome candidate 1 (WHSC1) (2). *NSD2* was first described as a gene dysregulated by the t(4;14)(p16.3;q32.3) translocation in approximately 15% of multiple myeloma (MM) cases and called Multiple Myeloma SET domain (MMSET) (2). (2-4). The translocation results in a fusion transcript of *NSD2* with the immunoglobulin heavy chain and increased NSD2 expression. The t(4;14) translocation can cause overexpression of both NSD2 and fibroblast growth factor receptor 3 (FGFR3) (2, 3). However, NSD2 is thought to be the primary oncogenic driver of the t(4;14)^+^ MM subtype because NSD2 is universally overexpressed, whereas the FGFR3 is not expressed in approximately 30% of MM cases (4–6). The role of NSD2 in driving t(4;14)^+^ MM pathogenesis was supported by knock-down of NSD2 in MM t(4;14)^+^ cell lines, which led to reduced growth and tumorigenesis (7–11). Conversely, transfection of t(4;14)^−^ cells with NSD2 promotes tumorigenesis and oncogenic transformation of primary cells via elevated levels of dimethylated H3K36 (H3K36me2) (12). Numerous studies have linked increased expression of NSD2 with globally increased levels of H3K36me2 (9, 12–21). High expression of NSD2 protein has been demonstrated in many different human cancer types, including bladder, brain, gastrointestinal, lung, liver, ovary, skin and uterus (18, 20, 22–28).

Notably, NSD2 is among the most frequently mutated genes in pediatric cancer genomes (29). The NSD2 SET domain mutation, E1099K, was identified in both acute lymphoblastic leukemia tumors and cell lines with increased H3K36me2 that lack the t(4;14) translocation (21, 30). Sequence results of >1000 pediatric cancer genomes, representing 21 different cancers, revealed the E1099K mutation in 14% of t(12;21) ETV6-RUNX1 containing ALLs (21). NSD2 is also among the most frequently mutated genes found in mantle cell lymphoma tumors, where both E1099K and T1150A mutations are observed (31). The E1099K mutation has also been reported in chronic lymphocytic leukemia (CLL), lung and stomach cancers (32–35). Recombinant NSD2 E1099K showed higher *in vitro* activity compared to the wild-type enzyme (21). Ectopic expression of NSD2 E1099K induced H3K36me2 and promoted transformation, while knockdown of the mutant enzyme reduced cell line proliferation and tumorigenesis (21).

Although NSD2 is an attractive therapeutic target, few small molecule inhibitors have been reported and none demonstrate the desirable attributes of high-quality chemical probes (36). The compound LEM-06 (IC_50_ = 900 μM) was discovered by virtual screening against an NSD2 homology model (37). The antiparasitic drug suramin inhibits NSD2 (IC_50_ = 0.3 - 21 μM), but is a pan-inhibitor of methyltransferases (38, 39) as well as other enzymes (40). Likewise, the nonspecific histone lysine methyltransferase inhibitor chaetocin (IC_50_ = 3 - 6 μM) showed similar inhibition of NSD1-3 (39). The natural product sinefungin is a close structural analogue of SAM and a modest inhibitor of NSD2 (IC_50_ = 26 - 30 μM) (41, 42). Structure-activity relationships have been reported for sinefungin analogues, the most potent of which inhibited the SET domains of NSD2 (IC_50_ = 1.8 μM) and SETD2 (IC_50_ = 0.29 μM) (41).

A major challenge in screening for small molecule inhibitors is that native NSD2 requires nucleosomes as a substrate (17). Recombinant NSD2 does not act on peptides and is thus not amenable to the commonly adapted histone-derived peptide screening platforms. Here we report biochemical assay development and pilot-scale screening using full-length recombinant wild-type, E1099K, and T1150A NSD2 enzymes with purified HeLa nucleosomes as a substrate. Chemical libraries were screened in 3-dose point quantitative high-throughput screening (qHTS) format (43) with the Methyltransferase-Glo (MTase-Glo™) methyltransferase bioluminescence assay (44). A number of counter and orthogonal assays were also developed to characterize the hits from the primary screen. Confirmed hits were validated with a radiolabeled SAM substrate-based NSD2 activity assay and further interrogated with binding studies by surface plasmon resonance (SPR), methyltransferase profiling, and activity assessments in U-2 OS cells.

## Results

### Primary assay development and optimization

To identify small molecule inhibitors of the wild-type NSD2, the MTase-Glo assay was adapted and optimized for use as a primary assay, with whole nucleosomes as the substrate, in 1,536-well format with 4 μL reaction volumes (Fig. 1B). The MTase-Glo assay reagent measures methyltransferase activity through the coupling of the NSD2 reaction product, S-adenosyl-L-homocysteine (SAH) to a bioluminescent signal (44). The assay was further optimized for the NSD2 mutants E1099K (Fig. 1C) and T1150A (Fig. 1D), to enable insight into cross-inhibition of clinically-relevant NSD2 enzymes. The enzyme concentrations were optimized to allow a robust signal-to-background ratio (>3.0) in a 15-min reaction at room temperature while consuming ≤20% substrate. The three optimized assays were robust with Z’-values near 0.9. There was no reduction in assay performance in the presence of up to 1.7% DMSO, which represents a triple aliquot of fixed volume (23 nL) transferred via a Kalypsys pintool equipped with a 1,536-pin array (Fig. 1E) (45). A titration of wild-type NSD2 demonstrated a linear correlation with methyltransferase activity as expected (Fig. 1F).

**Figure 1:**
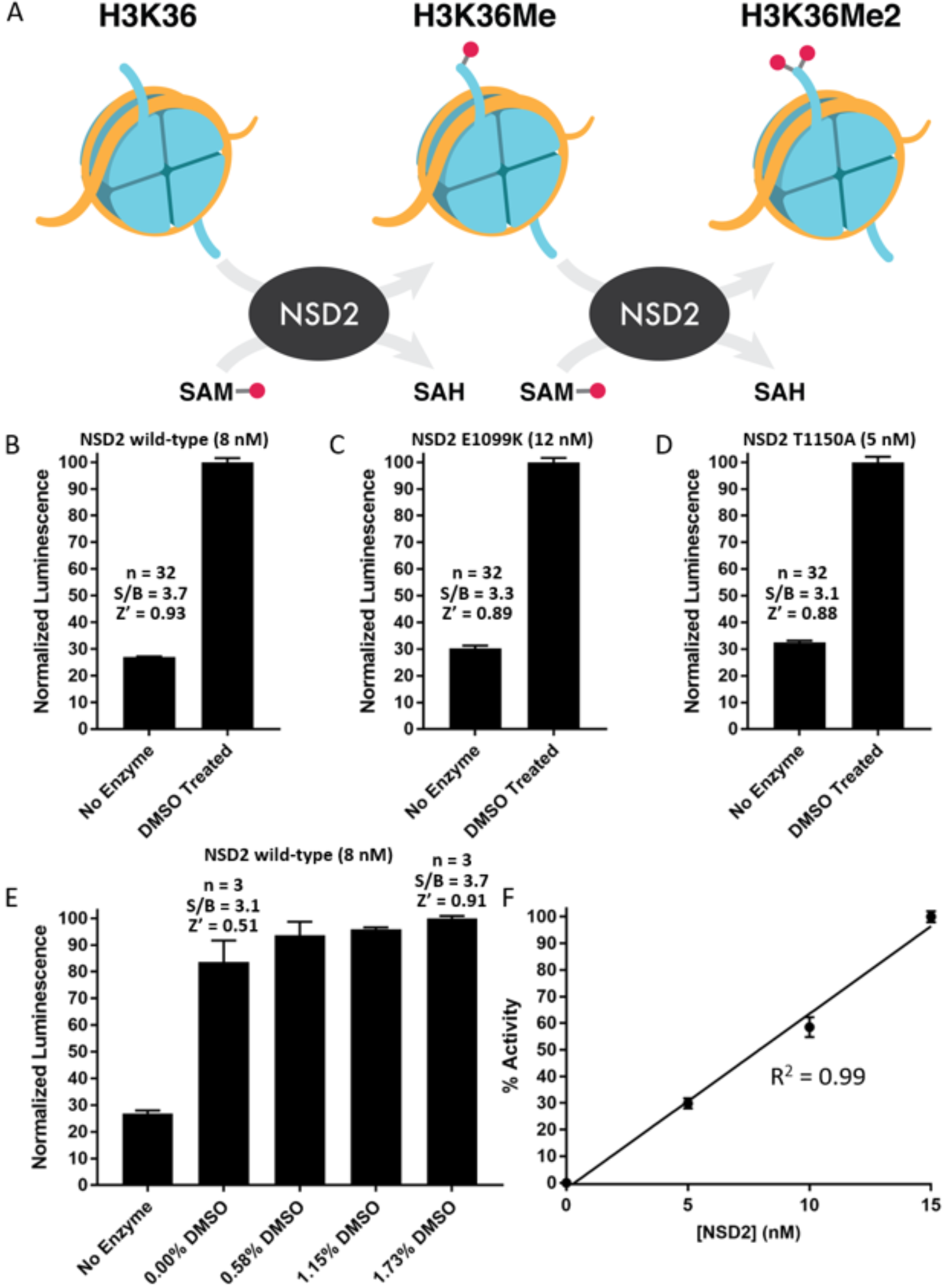
Optimization of the MTase-Glo primary assay. (A) Reaction scheme for the methylation of H3K36 by NSD2. (B-D) Optimized assay conditions for the full-length NSD2 wild-type (8 nM) (B) and mutant E1099K (12 nM) (C) or T1150A (5 nM) (D) enzymes with 500 nM nucleosomes, 0.58% DMSO and 1 μM SAM. The mean normalized luminescence values ± SD (*n* = 32) are plotted for B-D. The assays are robust with S/B values >3 and Z’-factor values ~0.9. (E) Titrations of DMSO demonstrated that the assay performance is not diminished by the introduction of vehicle up to 1.7% (mean ±SD; *n* = 3). (F) A linear correlation is observed between the wild-type NSD2 enzyme concentration and methyltransferase activity (mean ±SD; *n* = 8).

**Table 1.** Optimized protocol for the NSD2 MTase-Glo UltraGlo luciferase qHTS assay.

| Protocol Sequence for NSD2 MTase-Glo UltraGlo Luciferase qHTS Assay |  |  |  |
| --- | --- | --- | --- |
| Step | Parameter | Value | Description |
| 1 | Reagent | 4 μL | SAH standard curve in reaction buffer [1 μM - 0 μM] as a 7-point 1:2 dilution series ($n=4$), column 1 |
| 2 | Reagent | 3 μL | Nucleosome [666.7 nM] in reaction buffer, columns 2-3 |
| 3 | Reagent | 3 μL | NSD2 [10.7 nM] and nucleosome [666.7 nM] in reaction buffer, columns 4-48 |
| 4 | Controls | 23 nL | DMSO in columns 2-4 |
| 5 | Library Compounds | 23 nL | Columns 5-48 |
| 6 | Time | 30 min | Incubation |
| 7 | Reagent | 1 μL | S-adenosyl-L-methionine (SAM) [4 μM], columns 2-48 |
| 8 | Time | 15 min | Incubation |
| 9 | Reagent | 1 μL | MTase-Glo Reagent [5x] |
| 10 | Time | 30 min | Incubation |
| 11 | Reagent | 5 μL | MTase-Glo Detection Solution |
| 12 | Time | 30 min | Incubation |
| 13 | Detection | Luminescence | ViewLux uHTS Microplate Imager (PerkinElmer) |
| Step | Notes |  |  |
| 1,2,3 | White MB Greiner 1,536-well plates (789175-F); Reaction Buffer: 50 mM Tris-HCl, pH 8.8, 5 mM MgCl<sub>2</sub>, 50 mM NaCl, 1 mM TCEP and 0.01% Tween 20 |  |  |
| 4,5 | Pintool transfer (tip wash sequence: DMSO, MeOH, 3 sec vacuum dry) |  |  |
| 6, 8, 10 & 12 | Room temperature |  |  |
| 7 | Final reaction conditions (columns 4-48): 1 μM SAM, 500 nM nucleosome, 8 nM NSD2, 50 mM Tris-HCl, pH 8.8, 5 mM MgCl<sub>2</sub>, 50 mM NaCl, 1 mM TCEP and 0.01% Tween 20 | | |
| 9 | Conversion of SAH to ADP |  |  |
| 11 | Conversion of ADP to ATP and detection by UltraGlo luciferase |  |  |
| 13 | Settings: 30 sec exposure, 1x binning |  |  |

### Quantitative high-throughput screen (qHTS) for NSD2 inhibitors

The primary screen was carried out by performing qHTS against 16,251 compounds at 3 concentrations (115 μM, 57.5 μM and 11.5 μM) in 1,536-well plates (Table 1). Commercially-available libraries screened included the LOPAC (1,280 compounds), Prestwick (1,360 compounds), MicroSource (2,000 compounds), and Tocris (1,304 compounds) collections. Additionally, four NCATS libraries were screened, including an epigenetics-focused collection (284 compounds), a natural products library (2,108 compounds), the NPACT library (5,099 compounds) and the NCATS Pharmaceutical Collection (2,816 compounds) (46). These libraries, enriched with pharmacologically active compounds, were selected to evaluate the suitability of the primary and secondary assays to identify inhibitors of NSD2. The overall quality of the primary screen data from 43 plates was high, with average values for Z’-factor of 0.92 ± 0.02, signal-to-background (S/B) of 3.28 ± 0.31, and coefficient of variation (CV) for the DMSO control of 1.68% ± 0.49 (Fig. 2 A-C). The screen was conducted over a period of four months, so these data demonstrate excellent day-to-day reproducibility.

**Figure 2:**
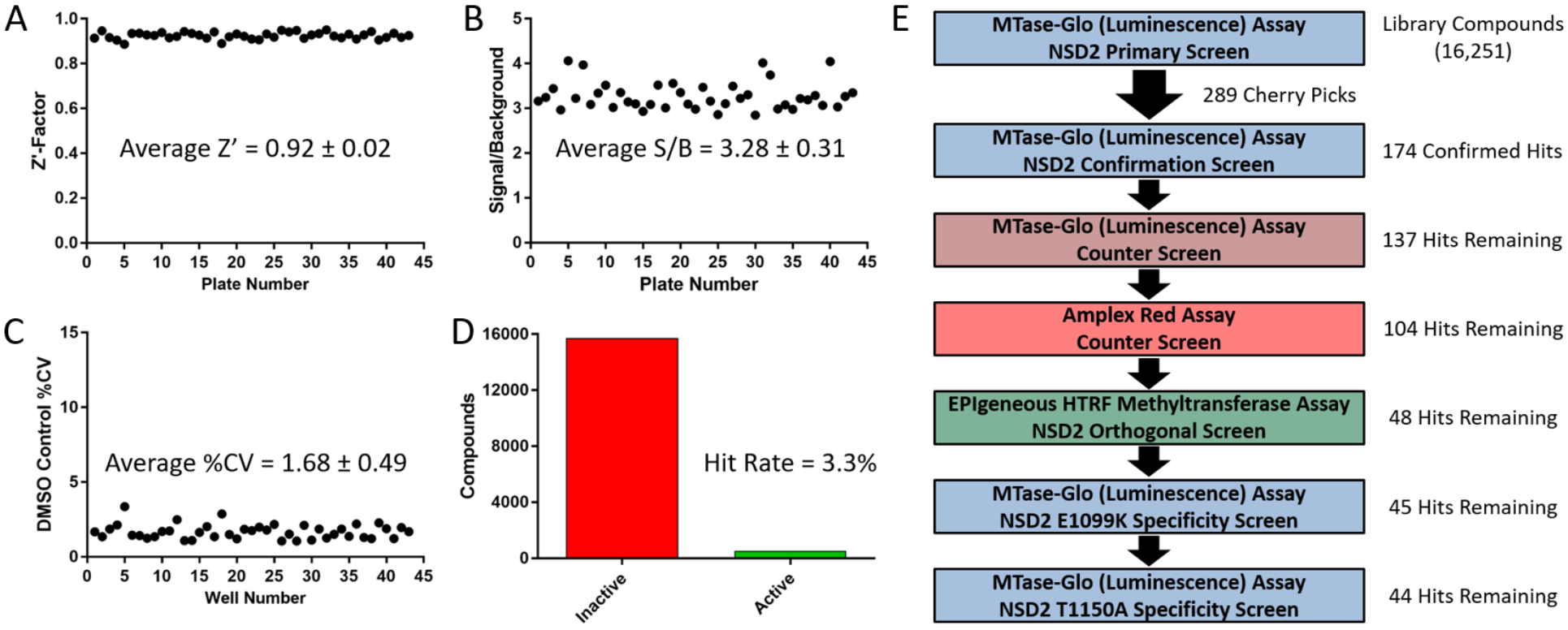
Quantitative high-throughput screening and secondary screening for inhibitors of wild-type and mutant NSD2 enzymes. (A-D) Statistics from the qHTS primary screen against full-length wild-type NSD2 with the MTase-Glo methyltransferase assay for forty-three 1,536-well plates. (A) The overall assay performance of the primary screen was robust with an average Z’-factor value of 0.92. (B) The average signal-to-background ratio was 3.28. (C) The average %CV for the DMSO control was 1.68. (D) Of the 16,251 compounds screened, 536 were identified as active (3.3% hit rate). (E) Primary and secondary assay triage process. Eight compound libraries (16,251 compounds) were screened in qHTS format at 3 concentrations, resulting in 536 hits. From the initial hit list, 289 compounds were selected as cherry picks and further evaluated in 11-point dose response. Of the 289 cherry picks, the activities of 48 were confirmed against NSD2 with the primary and orthogonal assays with no activity observed by the counter assays. The majority of these compounds were also active against the NSD2 mutants E1099K (45) and T1150A (44).

### Hit confirmation and secondary screening

Of the 16,251 compounds evaluated in the primary screen, 536 compounds were active with a maximum response ≥ 50%, corresponding to a hit rate of 3.3% (Fig. 2D). Promiscuity scores were calculated for each hit and any compounds with a promiscuity score higher than 0.2 were eliminated from the cherry-picking. Next the remaining hits were evaluated by a structural filter to eliminate the pan-assay interference compounds (PAINS) (47). Finally, 289 hit molecules were selected as cherry picks and prepared in 11-point concentration series from library stock solutions for further studies (Fig. 2E).

First, 174 of the cherry picks were confirmed as active in the primary assay (a 60% confirmation rate). Actives were defined as having concentration-response curves (CRCs) in the classes of 1.1, 1.2, 2.1, 2.2 and 3. In brief, classes 1.1 and 1.2 are the highest-confidence complete CRCs containing both upper and lower asymptotes with efficacies > 80% and ≤ 80%, respectively. Classes 2.1 and 2.2 are incomplete CRCs having only one asymptote with efficacies > 80% and < 80%, respectively. Class 3 CRCs show activity at only the highest concentration or are poorly fit. Class 4 CRCs are inactive, having a curve-fit of insufficient efficacy or lacking a fit altogether (43). Secondly, a MTase-Glo counter screen was implemented without NSD2 but containing 200 nM SAH (mimicking 20% substrate conversion), which identified 37 assay interference compounds that might act by inhibiting the coupling enzymes, luciferase or the luminescent signal. Potential redox cycling by compounds was assessed with an Amplex Red assay performed in the presence of reducing agents (48) and 63 compounds were found to be active with a threshold of 3σ. To provide additional evidence for on-target activity against NSD2, the orthogonal EPIgeneous Homogeneous Time-Resolved Fluorescence (HTRF) Methyltransferase Assay was utilized with reaction conditions identical to the primary assay (Fig. 3). The assay measures the NSD2 reaction product SAH, which competitively displaces d2-labeled SAH that is pre-bound to anti-SAH labeled with Lumi4-Tb, resulting in a loss of FRET signal (49).

**Figure 3:**
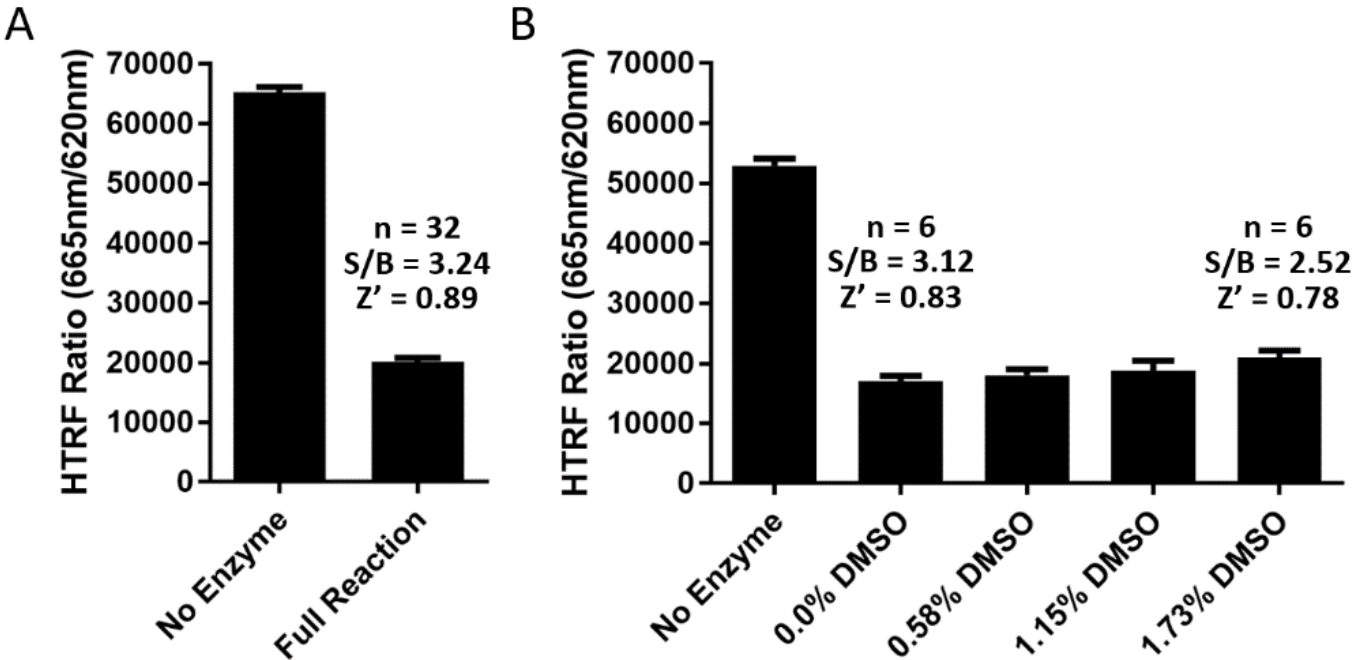
Optimized EPIgeneous Homogeneous Time-Resolved Fluorescence (HTRF) Methyltransferase Assay with the full-length NSD2 wild-type enzyme (8 nM), 500 nM nucleosomes, 0.58% DMSO and 1 μM SAM. (A) The mean HTRF ratio values ± SD for *n* = 32 replicates are plotted, showing robust performance with S/B values >3 and a Z’-factor value of ~0.9. (B) Titrations of DMSO demonstrated that the assay performance is not diminished by the introduction of vehicle (mean ±SD; *n* = 6).

Among the 37 compounds found to be active with the MTase-Glo counter assay was a known luciferase inhibitor NCGC00183809 (Fig. 4A). The compound shares structural commonality with the firefly luciferase inhibitor PTC-124 (44, 50, 51). As such it was likely a false-positive hit that inhibits the UltraGlo luciferase enzyme utilized in the MTase-Glo assay. This notion was supported by the results of the orthogonal assay, which did not indicate inhibition of NSD2 activity. UltraGlo luciferase is a genetically evolved firefly luciferase containing 70 mutations to improve its robustness, thermal stability, and resistance to interference compounds (52, 53), which is consistent with the 158-fold reduced potency of NCGC00183809 to UltraGlo compared with firefly luciferase (Fig. 4A). The nonspecific methyltransferase inhibitor DZNep (3-deazaneplanocin A) showed similar activities in both the MTase-Glo primary and counter assays; however, no inhibition of NSD2 activity was observed with the orthogonal assay (Fig. 4B).

**Figure 4:**
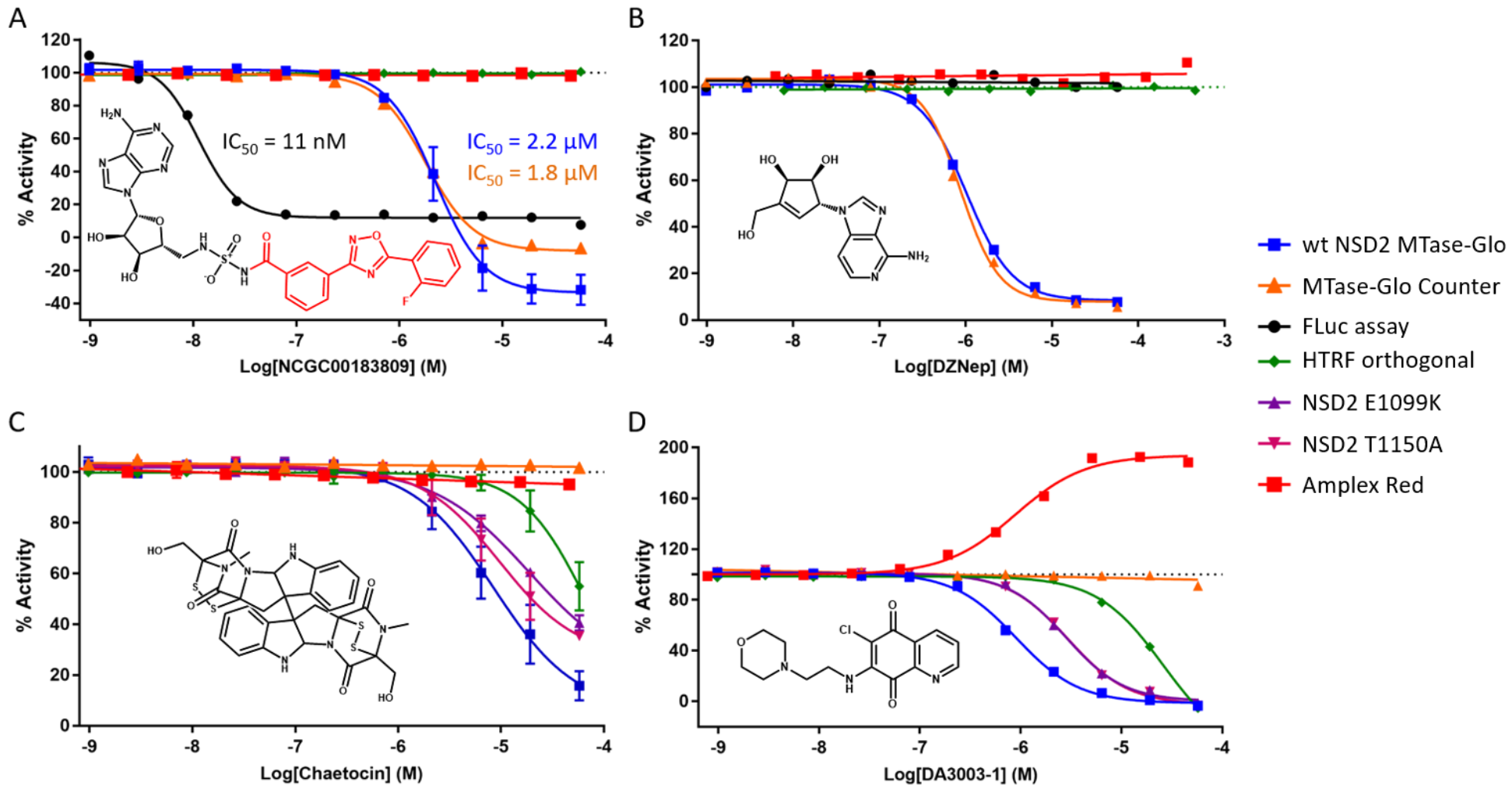
Primary and secondary assays characterize hit compounds against full-length wild-type and mutant NSD2 enzymes. (A) NCGC00183809 was identified by the primary assay (IC_50_ = 2.2 μM; Hill Slope = −1.84) and contains the PTC-124 moiety (colored red) (50, 51). The compound showed similar activity with the counter assay (IC_50_ = 1.8 μM; Hill Slope = −1.85), but was substantially more potent against firefly luciferase (IC_50_ = 11 nM; Hill Slope = −2.26). No activity against NSD2 was observed with the EPIgeneous HTRF orthogonal assay. The data of wild-type NSD2 MTase-Glo are shown as the mean value ±SD of *n* = 2 technical replicates. (B) DZNep showed similar activities between the primary (IC_50_ = 1 μM; Hill Slope = −1.63) and counter assays (IC_50_ = 0.87 μM; Hill Slope = −1.95). No activity was observed with the orthogonal assay. (C) Confirmation of chaetocin as an NSD2 inhibitor. Chaetocin was identified as a hit from the primary screen and the activity was confirmed by primary (IC_50_ = 8.5 μM; Hill Slope = −1.09) and orthogonal assays (IC_50_ = 67 μM; Hill Slope = −1.37). No activity was observed with the MTase-Glo counter assay or Amplex Red assay. Chaetocin also inhibits the NSD2 mutants E1099K (IC_50_ = 19 μM; Hill Slope = −0.93) and T1150A (IC_50_ = 9.6 μM; Hill Slope = −1.15). Data from the MTase-Glo Counter and Amplex Red assays resulted from single experiments. All other data are shown as mean values ±SD of *n* technical replicates (*n* = 3 for wild-type NSD2 MTase-Glo and *n* = 2 for HTRF orthogonal, NSD2 E1099K, and NSD2 T1150A). (D) The primary assay showed inhibition of wild-type (IC_50_ = 0.9 μM; Hill Slope = −1.4), E1099K (IC_50_ = 2.8 μM; Hill Slope = −1.51), and T1150A NSD2 (IC_50_ = 2.9 μM; Hill Slope = −1.44) enzymes by DA3003-1, while no activity was observed with the counter assay. Inhibition of wild-type NSD2 was further supported by the orthogonal assay. However, DA3003-1 showed activity with the Amplex Red assay (IC_50_ = 0.9 μM; Hill Slope = 1.25) indicating redox activity.

After the orthogonal assay, 48 confirmed hits remained that were not active in the two counter screens (Fig. 2E). Of the 48 confirmed inhibitors, 45 were also active against the E1099K enzyme, and 44 showed activity against the T1150A mutant. One of these hits is the nonspecific histone lysine methyltransferase inhibitor chaetocin, which was reported to inhibit NSD1-3 (39).

Chaetocin inhibited both wild-type and mutant NSD2 enzymes without showing activity by the MTase-Glo counter assay or Amplex Red assay (Fig 4C). The hit DA3003-1 is known to be redox active (48, 54), and this was corroborated by the Amplex Red counter screen (Fig 4D). Redox activity is undesirable, because it can result in nonspecific modulation of proteins, activation of cell pathways with redox-switches and cytotoxicity (55). DA3003-1 was nevertheless selected for follow-up studies due to its submicromolar potency with the primary assay (IC_50_ = 0.9 μM). In addition to chaetocin and DA3003-1, three other hits were selected for further studies that have not been previously linked to NSD2 inhibition: PF-03882845, TC LPA5 4, and ABT-199 (Table 2). In anticipation of a full-fledged HTS, these five compounds were selected to validate our post-HTS workflow, which is intended to further evaluate compound activities.

### Potency evaluation with the HotSpot radiolabel assay

Biochemical activities of the five selected hit compounds against wild-type and mutant NSD2 enzymes were further validated by the radioisotope-based HotSpot assay (Fig. 5). The HotSpot assay incorporates [^3^H]SAM to assess total histone methylation by direct measurement of the filter-bound tritiated substrate, without the need for coupling enzymes or antibodies (38). Both DA3003-1 and chaetocin inhibited wild-type and mutant NSD2 enzymes with submicromolar potencies, whereas PF-03882845, TC LPA5 4, and ABT-199 inhibited the enzymes at low-micromolar concentrations (Table 2). While chaetocin is known to inhibit NSD2, inhibition by DA3003-1, PF -03882845, TC LPA5 4 and ABT-199 has not been reported. Notably, the HotSpot assay was consistently more sensitive than the MTase-Glo and HTRF assays for both wild-type and mutant enzymes (Table 2). Overall the reaction conditions were very similar. The MTase-Glo and HTRF assays utilized 8 nM NSD2, 500 nM nucleosomes, 50mM Tris-HCl, pH 8.8, 5mM MgCl_2_, 50mM NaCl, 1mM TCEP, 0.01% Tween; whereas the HotSpot assay used 10 nM NSD2, 400 nM nucleosomes, 50 mM Tris-HCl, pH 8.5, 5 mM MgCl_2_, 50 mM NaCl, 1 mM DTT, 0.01% Brij35. A comparison of IC_50_ values determined by the HotSpot assay for all five compounds from reactions containing either 1 mM TCEP or DTT did not suggest that potency differences were due to the reducing agent (data not shown). Each of the five compounds inhibited wild-type, E1099K, and T1150A NSD2 enzymes with similar potencies (Table 2).

**Figure 5:**
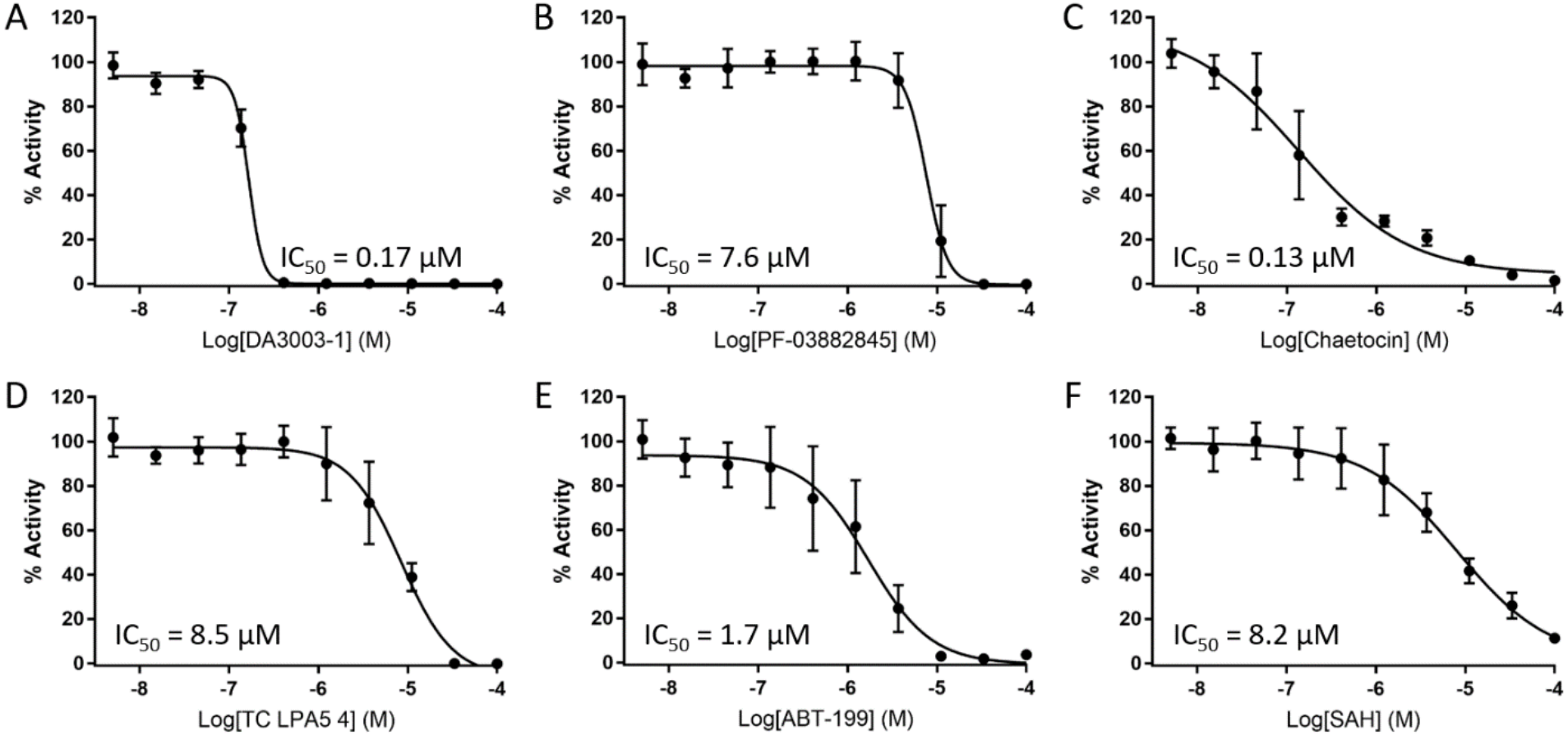
Inhibition of full-length wild-type NSD2 activity towards nucleosomes measured by the radiolabeled HotSpot assay with a 10-point concentration series of inhibitor. (A) DA3003-1 potently inhibits NSD2 with an IC_50_ value of 0.17 μM (Hill Slope = −5.57; *n* = 4), (B) PF-03882845 inhibits NSD2 activity with an IC_50_ value of 7.6 μM (Hill Slope = −3.68; *n* = 5), (C) Chaetocin is a potent inhibitor of NSD2 activity (IC_50_ = 0.13 μM; Hill Slope = −0.71; *n* = 5), (D) TC LPA5 4 inhibits NSD2 with an IC_50_ value of 8.5 μM (Hill Slope = −1.5; *n* = 5), (E) ABT-199 inhibits NSD2 with an IC_50_ value of 1.7 μM (Hill Slope = −1.23; *n* = 5), (F) The positive control SAH inhibits NSD2 with an IC_50_ value of 8.2 μM (Hill Slope = −0.85; *n* = 4). Data are plotted as the mean value ±SD of *n* technical replicate experiments.

**Table 2.**
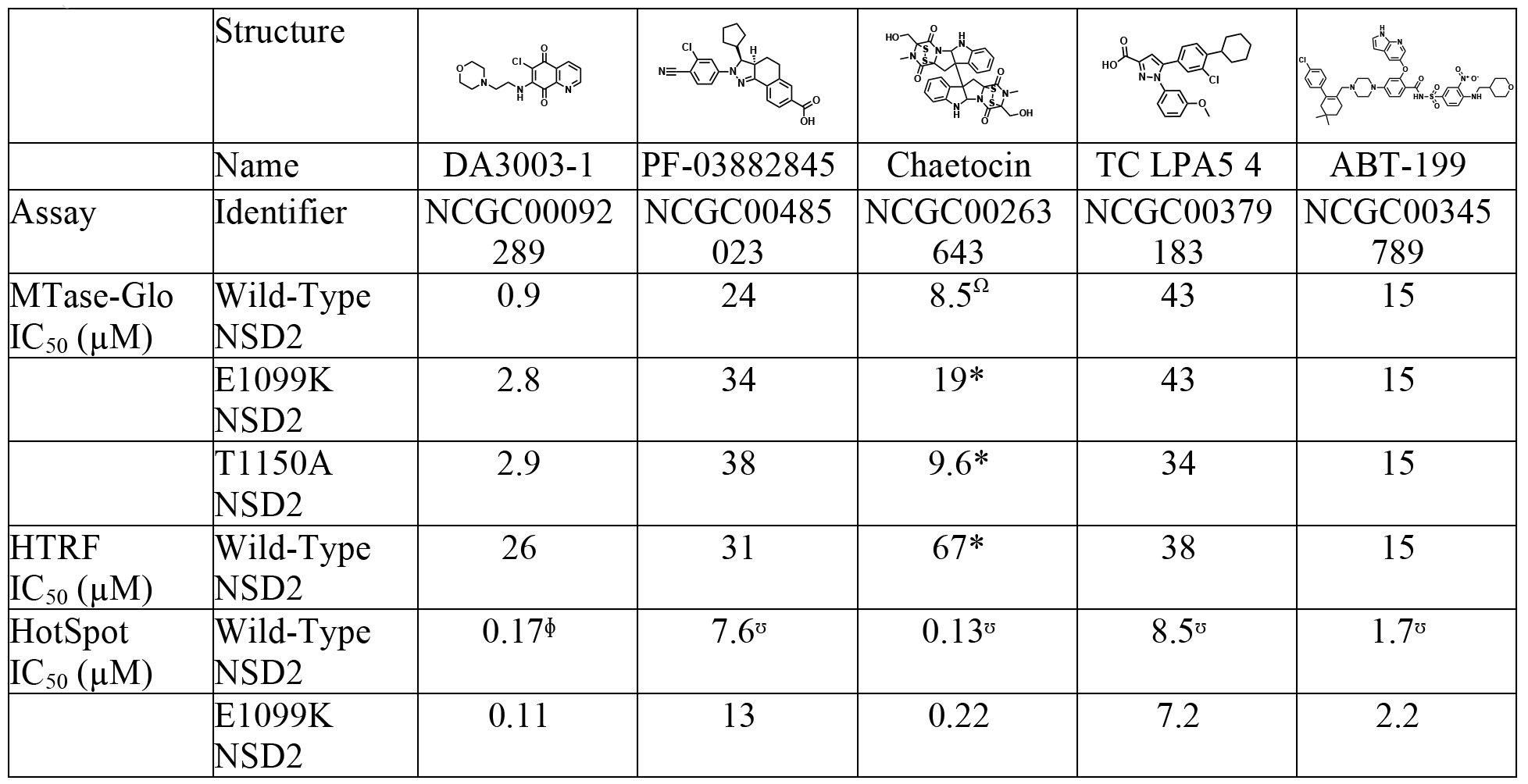
Biochemical activities of five compounds that inhibit wild-type and mutant NSD2 enzymes.

**Table 1.**
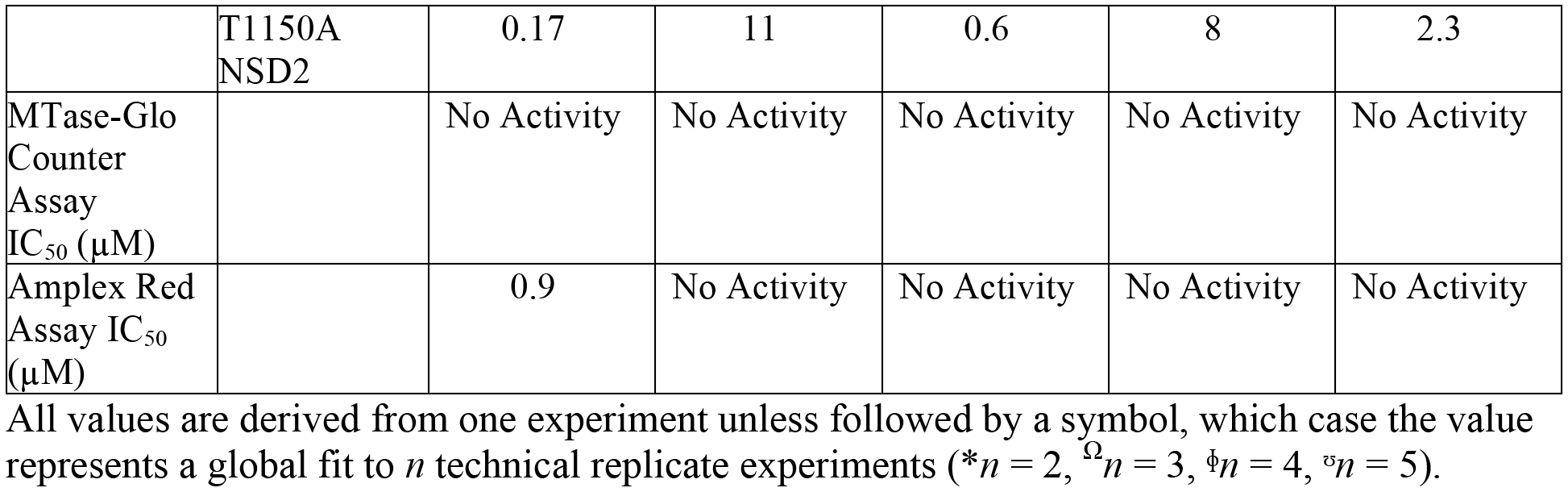

### Direct binding of inhibitors to the NSD2 SET domain

To support an on-target mechanism of action for the five inhibitors, surface plasmon resonance (SPR) was used to determine whether each inhibitor interacts with the catalytic SET domain of NSD2. As expected the two positive controls, cofactor SAM and product SAH, both bound the NSD2 SET domain with low micromolar affinities (Table 3). With the exception of chaetocin, the inhibitors bound the NSD2 SET domain stoichiometrically, with dissociation constants (*K*_d_) comparable to the *in vitro* IC_50_ values determined by the HotSpot assay (Table 2). Chaetocin bound the SET domain with an apparent dissociation constant of 20 nM. Additionally, the data indicated super-stoichiometric binding, which might be due to a binding ratio higher than 1:1 that is consistent with chaetocin’s two disulfide moieties forming adducts with the NSD2 protein, or compound aggregation (data not shown) (56). These data indicate that all five compounds might mediate inhibition of NSD2 by directly binding to the catalytic SET domain.

**Table 3.** Direct binding of hit compounds to the NSD2 SET domain as measured by surface plasmon resonance, with SAH and SAM as positive controls.

| Compound Name | $k_{on}$ (M<sup>-1</sup>s<sup>-1</sup>) | $k_{off}$ (s<sup>-1</sup>) | $K_d$ (μM) |
| --- | --- | --- | --- |
| SAH | $2.33 \times 10^3$ | $1.19 \times 10^{-2}$ | 5.1 |
| SAM | $4.06 \times 10^3$ | $1.70 \times 10^{-2}$ | 4.2 |
| DA03003-1 | $3.67 \times 10^4$ | $1.34 \times 10^{-2}$ | 0.37 |
| PF-03882845 | $8.55 \times 10^3$ | $3.31 \times 10^{-2}$ | 3.9 |
| Chaetocin | $2.21 \times 10^3$ | $4.59 \times 10^{-5}$ | 0.02 |
| TC LPA5 4 | $4.29 \times 10^3$ | $3.59 \times 10^{-2}$ | 8.4 |
| ABT-199 | $4.39 \times 10^3$ | $3.64 \times 10^{-2}$ | 8.3 |

### Selectivity assessment by methyltransferase profiling

Methyltransferase profiling of the five NSD2 inhibitors was performed to examine the selectivity towards NSD2 compared to other methyltransferases. Activities of the inhibitors were evaluated against 35 other methyltransferases, including NSD1 and NSD3, with the HotSpot methyltransferase assay technology (Table 4). DA3003-1 was the least selective methyltransferase inhibitor and showed potent inhibition of nearly every enzyme except DOT1L (no activity) and GLP (weak inhibition). Furthermore, DA3003-1 inhibited 27 enzymes potently with submicromolar IC_50_ values. Both PF-03882845 and TC LPA5 4 inhibited most enzymes, albeit with weak potencies, but showed some selectivity to the PRMT5/MEP50 complex and the MLL4 complex. Chaetocin inhibited 18 methyltransferases, twelve with submicromolar potencies (including all NSD enzymes) and two with IC_50_ values above 75 μM. ABT-199 inhibited 23 methyltransferases, two of which have IC_50_ values above 65 μM. The NSDs were among the most potently inhibited enzymes, with IC_50_ values in the low-micromolar range.

**Table 4.** Selectivity of inhibitor compounds towards wild-type NSD2 was assessed by profiling 37 other methyltransferases, including the E1099K and T1150A NSD2 mutants, with the HotSpot methyltransferase assay. The IC_50_ values for all test compounds are colored as a heat map with green indicating potent inhibition and red indicating weak inhibition. No color indicates no measurable activity. All values are derived from a single experiment except for those of wild-type NSD2, which result from a global fit of multiple data sets as indicated in Table 2.

| Methyltransferase | Substrate | DA3003-1<br>IC<sub>50</sub> (μM) | PF-03882845<br>IC<sub>50</sub> (μM) | Chaetocin<br>IC<sub>50</sub> (μM) | TC LPA5 4<br>IC<sub>50</sub> (μM) | ABT-199<br>IC<sub>50</sub> (μM) | Control<br>IC<sub>50</sub> (μM) | Control ID |
| --- | --- | --- | --- | --- | --- | --- | --- | --- |
| ASH1L | Nucleosomes | 0.2 | 9.8 | 0.31 | 9.2 | 3.3 | 0.1 | Chaetocin |
| DNMT1 | Poly (dl-dC) | 1.7 | 14 | No Inhibition | 12 | 44 | 0.13 | SAH |
| DNMT3a | Lambda DNA | 1.1 | 18 | No Inhibition | 16 | 18 | 0.12 | SAH |
| DNMT3b | Lambda DNA | 0.38 | 19 | No Inhibition | 24 | >100.00 | 0.065 | SAH |
| DNMT3b/3L | Lambda DNA | 0.72 | 31 | No Inhibition | 30 | No Inhibition | 0.012 | SAH |
| DOT1L | Nucleosomes | No Inhibition | 14 | No Inhibition | 8.2 | 2 | 0.74 | SAH |
| EZH1 Complex | Core Histone | 0.37 | 4 | No Inhibition | 40 | 66 | 19 | SAH |
| EZH2 Complex | Core Histone | 0.46 | 3.8 | No Inhibition | 40 | 23 | 17 | SAH |
| EZH2 (Y641F) Complex | Core Histone | 0.68 | 3.8 | No Inhibition | 31 | 21 | 68 | SAH |
| G9a | Histone H3 (1-21) | 0.9 | No Inhibition | 2.6 | No Inhibition | No Inhibition | 0.71 | SAH |
| GLP | Histone H3 (1-21) | 36 | No Inhibition | 77 | No Inhibition | No Inhibition | 0.53 | SAH |
| MLL1 Complex | Nucleosomes | 0.25 | 6.1 | 0.32 | 3.2 | 0.72 | 0.55 | SAH |
| MLL2 Complex | Nucleosomes | 0.32 | 15 | 3 | 18 | 4.6 | 27 | SAH |
| MLL3 Complex | Core Histone | 1.1 | 29 | 1.2 | 63 | No Inhibition | 57 | SAH |
| MLL4 Complex | Nucleosomes | 0.22 | 1.3 | 0.07 | 1 | 0.52 | 2.5 | SAH |
| NSD1 | Nucleosomes | 0.11 | 12 | 0.06 | 12 | 2.4 | 9.7 | SAH |
| NSD2 | Nucleosomes | 0.17 | 7.6 | 0.13 | 8.5 | 1.7 | 6.2 | SAH |
| NSD2 (E1099K) | Nucleosomes | 0.11 | 13 | 0.22 | 7.2 | 2.2 | 0.94 | SAH |
| NSD2 (T1150A) | Nucleosomes | 0.17 | 11 | 0.6 | 8 | 2.3 | 1.5 | SAH |
| NSD3 | Nucleosomes | 0.26 | 3.2 | 0.33 | 2.3 | 1.3 | 0.09 | Chaetocin |
| PRDM9 | Histone H3 | 0.37 | 85 | 5.9 | No Inhibition | No Inhibition | 8.1 | Chaetocin |
| PRMT1 | Histone H4 | 0.26 | 8.5 | No Inhibition | 68 | 83 | 0.17 | SAH |
| PRMT3 | Histone H4 | 0.31 | 29 | No Inhibition | 5.7 | 36 | 2.3 | SAH |
| PRMT4 | Histone H3 | 1.4 | 29 | No Inhibition | No Inhibition | No Inhibition | 0.29 | SAH |
| PRMT5 | Histone H2A | 0.65 | 11 | No Inhibition | 17 | No Inhibition | 49 | SAH |
| PRMT5/MEP50 Complex | Histone H2A | 0.21 | 0.38 | No Inhibition | 0.33 | No Inhibition | 1 | SAH |
| PRMT6 | GST-GAR | 1 | 70 | No Inhibition | No Inhibition | No Inhibition | 0.084 | SAH |
| PRMT7 | GST-GAR | 0.42 | No Inhibition | 156 | No Inhibition | No Inhibition | 0.045 | SAH |
| PRMT8 | Histone H4 | 0.22 | 21 | No Inhibition | 71 | 34 | 0.063 | SAH |
| SET1b Complex | Core Histone | 0.26 | 4.3 | 0.19 | 8 | 7.1 | 1.5 | SAH |
| SET7/9 | Core Histone | 1.4 | 31 | No Inhibition | 52 | No Inhibition | 83 | SAH |
| SET8 | Nucleosomes | 4.4 | 42 | No Inhibition | 44 | 5.3 | 0.95 | Ryuvidine |
| SETD2 | Nucleosomes | 0.21 | 13 | 0.45 | 13 | 3.3 | 5.2 | SAH |
| SMYD1 | Core Histone | 1 | 11 | No Inhibition | 21 | No Inhibition | 37 | SAH |
| SMYD2 | Histone H4 | 1.7 | 32 | No Inhibition | 21 | 44 | 4.5 | LLY 507 |
| SUV39H1 | Histone H3 | 0.47 | No Inhibition | 0.74 | No Inhibition | No Inhibition | 25 | SAH |
| SUV39H2 | Histone H3 | 0.62 | No Inhibition | 0.57 | No Inhibition | No Inhibition | 55 | SAH |
| SUV420H1TV2 | Nucleosomes | 0.34 | 4.4 | No Inhibition | 4.2 | 2.2 | 31 | SAH |

### Inhibitor activities in a cell-based assay

To evaluate the activities of the five NSD2 inhibitors in cells, U-2 OS human osteosarcoma cells were chosen due to a relatively high expression of endogenous NSD2 protein (57). The cells were treated with each of the five compounds in 10-point dose response from 0.0025 - 50 μM for 96 hours. NSD2 inhibition should result in reduced H3K36 di-methylation. Total histone H3 and H3K36me2 levels were measured by western analysis. The positive control DZNep was tested at a concentration of 10 μM in parallel with each compound and also evaluated in dose-response (Fig. 6A). Densitometry was used to quantify both H3K36me2 and total H3 and the densities of H3K36me2 were normalized to those of H3 (Fig. 6B). The growth of U-2 OS cells over 96 hours in the presence of test compounds at the same concentrations was also evaluated (Fig. 6C). The IC_50_ value of DZNep for reducing H3K36me2 levels in U-2 OS cells was 390 nM with a modest dose-dependent reduction in U-2 OS confluency (AC_50_ = 180 nM). Similar to DZNep, DA3003-1 treatment also resulted in a dose-dependent reduction in H3K36me2; however, higher drug concentrations resulted in cytotoxicity (CC_50_ = 270 nM) similar to chaetocin and the control bortezomib, both of which were cytotoxic at all concentrations tested. PF-03882845 reduced H3K36me2 (IC_50_ = 3.2 μM) over a range of concentrations that had negligible influence on growth; however, significant cell death at concentrations above 16.7 μM was observed during the western analysis experiment (Fig. 6A), so data from the two highest concentrations were excluded from the potency calculation. TC LPA5 4 did not reduce H3K36me2 over the concentrations tested and minimal reduction in growth was observed. ABT-199 did not appear to reduce H3K36me2 levels, however, concentrations above 17 μM were cytotoxic.

**Figure 6:**
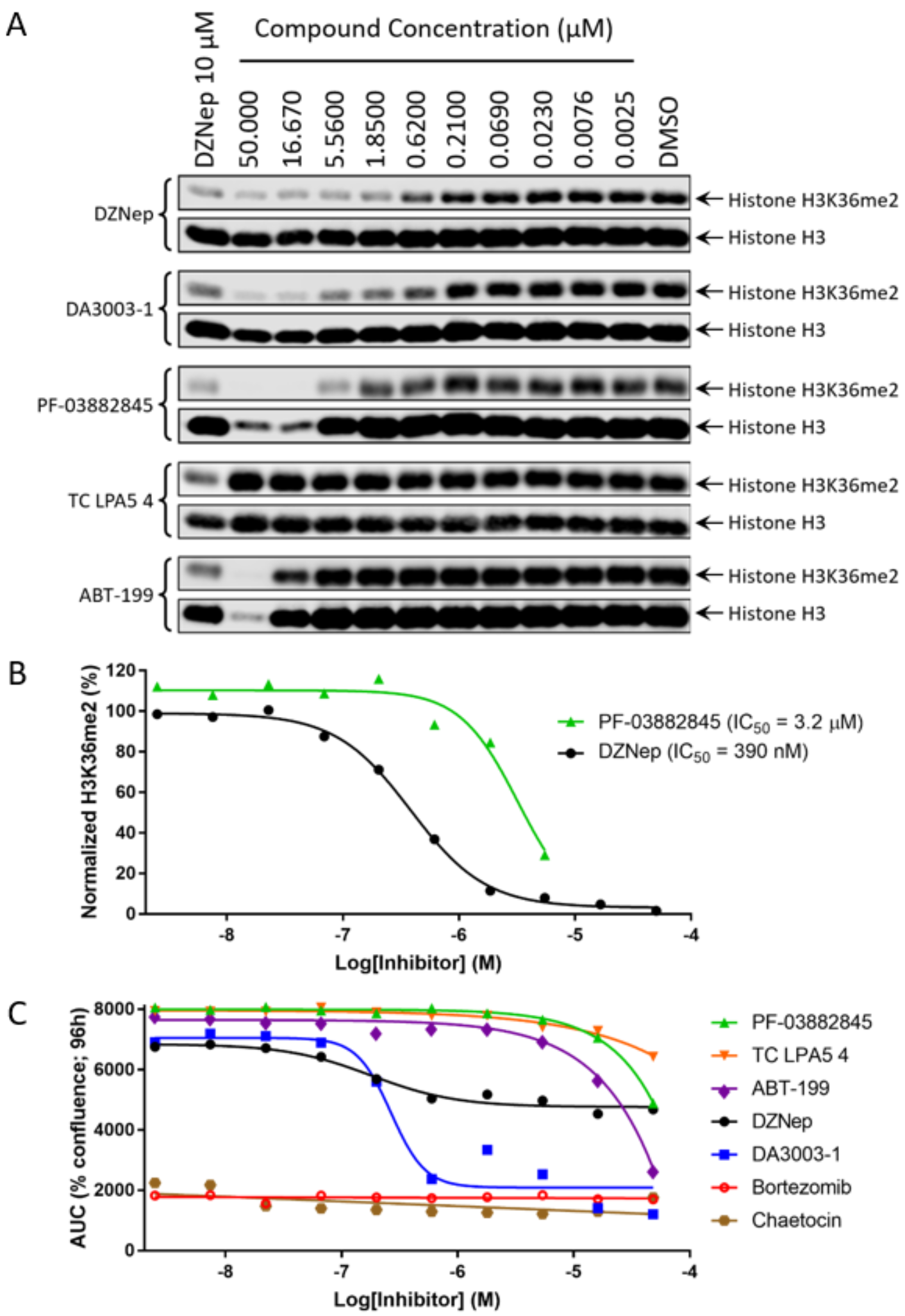
Analysis of inhibitor activities towards U-2 OS human osteosarcoma cells. (A) U-2 OS cells were treated with the indicated concentrations of test compounds or 10 μM of the global histone methyltransferase inhibitor DZNep, which served as a positive control. After 96 hours of treatment, whole cell lysates were subjected to western blot analyses. The blots were probed with anti-histone H3K36me2 and anti-histone H3 antibodies, respectively. (B) Densitometry was performed and the values for H3K36me2 densities were normalized to the total H3 densities. The resulting data for PF-038828455 (IC_50_ = 3.2 μM; Hill Slope = −1.74) and DZNep (IC_50_ = 390 nM; Hill Slope = −1.34) were plotted and fit to a nonlinear four parameter equation. (C) U-2 OS cells were treated with the indicated concentrations of test compounds and percent confluency of the cells at 96 hours is shown as area under the curve (AUC). Bortezomib was included as a control for cytotoxicity.

## Discussion

A number of histone lysine methyltransferases have been implicated as attractive therapeutic targets in the field of oncology and small molecule inhibitors are in different stages of preclinical and clinical development (58, 59). Selective inhibitors of NSD2 are of major interest both to advance basic research and for therapeutic development. However, NSD2 is regarded as a challenging target (60) and no selective inhibitors of NSD2 have been reported to date. Challenges in studying NSD2 *in vitro* include that the target of NSD2 methylation depends on the nature of the substrate (17) and full-length NSD2 is only active against a nucleosome substrate (38), which may be cost prohibitive in many cases.

A wide variety of methyltransferase assays have been described in the literature (38, 44, 61–66); however, few optimized assays have been reported for NSD2 in plate formats to enable HTS. Recently a radiolabeled [^3^H]SAM assay was optimized for the NSD2 SET domain with both histone octamer (Z’ = 0.69) and nucleosome (Z’ = 0.75) substrates in a 384-well format (39). The assay utilizing a nucleosome substrate was used to screen 1,040 compounds from the Prestwick Chemical Library at a single concentration of 25 μM. While the reported assays show robust performance in 384-well format, the use of radiolabeled reagents for HTS is a challenge for many laboratories due to safety regulations and disposal costs.

Herein we report the implementation and validation of two optimized homogenous NSD2 activity assays in the highly miniaturized 1,536-well format for the identification of small molecule inhibitors from chemical libraries. The assays were utilized to screen the full-length wild-type NSD2 enzyme against a nucleosome substrate in qHTS format with three concentrations of test compounds. The use of qHTS reduces both false positive and false negative hits common to single-point HTS and facilitates selection of actives (43). For the primary screen we used the recently reported Methyltransferase-Glo assay reagent with a sensitive bioluminescent readout (44). A similar approach has been applied for the discovery of NSD1 inhibitors by HTS (67). After screening eight libraries, including numerous pharmacologically active collections, containing over 16,000 compounds, many hits were identified including chaetocin, which is known to inhibit NSD2 (39). By incorporating orthogonal and counter screens hits were prioritized for subsequent follow-up studies.

Among the confirmed active inhibitors, DA3003-1, PF-03882845, chaetocin, TC LPA5 4, and ABT-199 were selected for further characterization. *In vitro* potencies were determined by the HotSpot assay, which is a direct readout of the NSD2 reaction product (38). The HotSpot assay is very similar in format to the traditional gold standard radioisotope detection used in conjunction with gel electrophoresis or mass spectroscopy. The five compounds inhibited wild-type NSD2 as well as the E1099K and T1150A mutant enzymes.

DA3003-1 is a cell-permeable Cdc25 phosphatase inhibitor that potently and irreversibly inhibits all Cdc25 isoforms, including Cdc25A (IC_50_ = 29 nM), Cdc25B2 (IC_50_ = 95 nM), and Cdc25C (IC_50_ = 89 nM) (68). It is known that DA3003-1 is capable of redox cycling (48, 54), which was verified here by the Amplex Red counter screen. In addition to potent inhibition of NSD2 activity, our data demonstrates that DA3003-1 bound the SET domain with a strong affinity (*K*_d_ = 370 nM, Table 3) which is comparable to its potency (IC_50_ = 170 nM, Table 2). Together this suggests that DA300-1 inhibits NSD2 through a direct interaction with the catalytic SET domain, although it is most likely nonspecific. Notably, DA3003-1 inhibited 27 of the 38 methyltransferase enzymes tested with submicromolar potencies and another eight with IC_50_ values near 1 μM. The potency of DA3003-1 in cells with respect to reducing H3K36me2 correlates with cytotoxicity (CC_50_ = 270 nM).

The Pfizer compound PF-03882845 is a highly potent mineralocorticoid receptor antagonist (IC_50_ = 0.755 nM) (69). In comparison, it inhibited NSD2 *in vitro* with an approximately 10,000-fold weaker potency (IC_50_ = 7.6 μM) and bound the SET domain (*K*_d_ = 3.9 μM) within 2-fold of the IC_50_ value. Profiling studies indicated that PF-03882845 inhibits many other methyltransferases with modest potencies, though it inhibits the histone arginine methyltransferase PRMT5 with a submicromolar potency. Interestingly, PF-03882845 reduced H3K36me2 in cells with a potency (3.2 μM) within 2-fold of the biochemical IC_50_ value. Concentrations at and above 17 μM resulted in substantial cell death in the western analysis experiment, but only minimal reductions in cell growth were observed with up to 50 μM drug at 96 hours in the cell growth assay.

TC LPA5 4 was first reported by Sanofi Aventis as a selective lysophosphatidic acid receptor 5 (LPA_5_) antagonist that inhibited LPA-mediated human platelet aggregation with an IC_50_ value of 2.2 μM (70). The biochemical potency of TC LPA5 4 against NSD2 (IC_50_ = 8.5 μM) was nearly identical to its affinity to the catalytic SET domain (*K*_d_ = 8.4 μM). The methyltransferase activity profiles of TC LPA5 4 and PF-03882845 showed striking similarities (Table 4). Both compounds inhibited wild-type NSD2 with an IC_50_ value near 8 μM. Also, both compounds were most potent against the PRMT5/MEP50 complex with nearly identical IC_50_ values. Furthermore, both compounds had approximately 3-fold weaker potency against the MLL4 complex. Compared to the PRMT5/MEP50 complex, the potency values of PF-03882845 were at least 8-fold weaker against 31 other methyltransferases. Similarly, compared to the PRMT5/MEP50 complex, the potency values of TC LPA5 4 is at least 7-fold weaker against 28 other methyltransferases. No activity or toxicity was observed for TC LPA5 4 in U-2 OS cells.

The fungal mycotoxin chaetocin is known to inhibit NSD2 (39), so identifying it as a hit further validated our screening approach. Of the 38 methyltransferases profiled, chaetocin inhibited 12 with submicromolar potencies. Notably the methyltransferase profiling indicated chaetocin potencies of 740 nM against SUV39H1 and 570 nM against SUV39H2, which is consistent with a previously reported value of 600 nM (71). Chaetocin was initially reported to be a specific inhibitor of the histone lysine methyltransferase SU(VAR)3-9 both *in vitro* and *in vivo* (71). However, the two disulfide bonds of chaetocin can complicate bioassay interpretation because of the potential for redox activity and covalent modification of proteins (36). Indeed reports have indicated that the activity against histone lysine methyltransferases is due to chemical modification of the enzyme by the disulfide groups (72, 73). Superstoichiometric binding of chaetocin to the NSD2 SET domain was observed by surface plasmon resonance, which might be due to the formation of direct compound-thiol adducts. The affinity of chaetocin to the NSD2 SET domain was strong (*K*_d_ = 20 nM) with a biochemical potency about 7-fold weaker. Chaetocin was cytotoxic to U-2 OS cells at all concentrations tested.

ABT-199, also known as GDC-0199 or venetoclax, binds BCL-2 with a subnanomolar affinity (*K*_*i*_ < 0.010 nM) and is approved by the FDA for the treatment of CLL (74, 75). The biochemical potency of ABT-199 against wild-type NSD2 (IC_50_ = 1.7 μM) was similar against the NSD2 mutants, NSD1 and NSD3 (Table 4). The compound was most potent against the MLL1 and MLL4 complexes. The affinity of ABT-199 to the NSD2 SET domain (*K*_d_ = 8.3 μM) is nearly 5-fold weaker than the biochemical potency. At a concentration of 50 μM, ABT-199 was cytotoxic, consistent with its use in oncology, but any influences on cellular H3K36me2 levels were negligible.

DZNep is a carbocyclic analog of adenosine and a derivative of the antibiotic neplanocin-A. It was initially reported as a competitive inhibitor of S-adenosyl homocysteine hydrolase at picomolar concentrations (76). DZNep has since been reported as a global histone methylation inhibitor when used at substantially higher concentrations (36, 77, 78). Although DZNep was identified as a hit from the primary screen, similar activities were observed with the MTase-Glo primary and counter assays and no inhibition of NSD2 activity was observed with the orthogonal assay. The potency of DZNep in reducing H3K36me2 in U-2 OS cells (IC_50_ = 390 nM; Fig. 6) is similar to its potency in reducing H3K27me3 in SU-DHL-6 cells (IC_50_ = 160 nM) when assessed by the same method of western analysis (79). DZNep has been also shown to reduce H3K36me2 in SW480 cells at a concentration of 5 μM (80).

The purpose of this study was to establish a HTS discovery pipeline for NSD2, and to evaluate the workflow for identifying high-quality tool inhibitors of NSD2. The majority of the molecules screened were from pharmacologically active libraries that served to validate the primary and secondary assays. The identification of known methyltransferase inhibitors, including chaetocin and DZNep, further validated the workflow. During the course of this pilot, many known interference compounds were identified by the secondary assays, thereby demonstrating how such bad actors behave among the various assays. Five actives selected from the primary screen were shown to bind the catalytic SET domain and inhibit NSD2 activity *in vitro*. Although these studies confirm inhibition of NSD2, they do not rule out inhibition by intractable mechanisms of action, such as non-specific reactivity, redox, or aggregation. Two of the five compounds reduced H3K36me2 in U-2 OS cells, but the mechanisms are likely to be complicated and involving multiple targets. These studies provide a basis for the future discovery and development of novel selective NSD2 inhibitors by establishing a robust workflow for identifying and triaging hits from high-throughput screens.

## Experimental Procedures

### Chemicals, Reagents and Libraries

MTase-Glo (V7602) was purchased from Promega, Inc. (Madison, WI). The EPIgeneous HTRF Methyltransferase Assay (Cat# 62SAHPEB) was purchased from Cisbio (Bedford, MA). Dimethyl sulfoxide (DMSO) was purchased from Fisher Scientific (Pittsburgh, PA). The libraries screened include the Library of 1,280 Pharmacologically Active Compounds (LOPAC^1,280^; Sigma-Aldrich, St. Louis, MO), Tocris (1,304 compounds, Tocris Bioscience, Bristol, United Kingdom), Prestwick (1,360 compounds, Prestwick Chemical, San Diego, CA), MicroSource (2,000 compounds, MicroSource Discovery Systems, Gaylordsville, CT), NPC (the National Center for Advancing Translational Sciences [NCATS] Pharmaceutical Collection (2,816 compounds) (46), NPACT (NCATS Pharmacologically Active Chemical Toolbox, 5,099 compounds), an epigenetic collection (284 compounds), and a natural products library (2,108 compounds). The reference compound DZNep (3-Deazaneplanocin A) was purchased from Selleckchem and dissolved with DMSO to a 10 mM stock. The U-2 OS human osteosarcoma cell line was purchased from American Type Culture Collection (Manassas, VA). U-2 OS cells were grown in McCoy’s 5A Medium supplemented with 10% FBS. 100 μg/ml penicillin and 100 μg/ml streptomycin were added to the culture media. Cultures were maintained at 37 °C in a humidified atmosphere of 5% CO_2_ and 95% air. 12% Bis-Tris gel and nitrocellulose membrane were purchased from Thermo Fisher Scientific. Di-Methyl-Histone H3 (Lys36) Rabbit mAb (#2901) and Histone H3 Mouse mAb (#3638) were purchased from Cell Signaling Technology. Anti-rabbit IgG IRDye 680RD and anti-mouse IgG IRDye 800CW secondary antibodies were purchased from LI-COR.

### Enzymes and Substrates

Human ASH1L (residues 2046-2330; accession # NM_018489) was expressed in Escherichia coli as N-terminal polyhistidine tag. Human DNMT1 (residues 2-1632; accession # NM_001130823) was expressed in an insect cell/baculovirus expression system as N-terminal GST fusions. Human DNMT3a (residues 2-912; accession # NM_175629) was expressed in an insect cell/baculovirus expression system as N-terminal GST fusions. Human DNMT3b (residues 2-853; accession # NM_006892) was expressed in an insect cell/baculovirus expression system as N-terminal GST fusions. Complex of Human DNMT3b (residues 564-853; accession # NM_006892) with N-terminal polyhistidine tag and Human DNMT3L (residues 160-387; accession # NM_013369) with N-terminal GST tag were coexpressed in an insect cell/baculovirus expression system. Human DOT1L (residues 1–416; accession # NM_032482) was expressed in Escherichia coli as N-terminal GST fusions. Human recombinant EZH1 (residues 2–747; Genbank Accession #NM_001991) or EZH2 (residues 2–746; Genbank Accession #NM_001203247) were coexpressed with human recombinants AEBP2 (2–517; NM_001114176), EED (2–441; NM_003797), RbAp48 (2–425; NM_005610), and SUZ12 (2–739; NM_015355) in an insect cell/baculovirus expression system to form the 5-member EZH1 or EZH2 complexes. All proteins were full length (residue 2 through C-terminus). The EED subunit incorporated an N-terminal Flag-tag, and all others included an N-terminal polyhistidine tag. Human GLP (residues 894–1298; accession # NM_024757) and Human G9a (residues 786–1210; accession # NM_006709) were expressed as N-terminal GST fusion protein in *E. coli*. Human MLL1 (residues 3745–3969; accession #NM_005933), human WDR5 (22–334; NM_017588), RbBP5 (1–538; NM_005057), Ash2L (2–534; NM_001105214), and DPY-30 (1–99; NM_0325742) were expressed in *E. coli* with N-terminal polyhistidine tag assembled as a complex (2 copies of DPY-30) and stored in 20mM Tris–HCl, pH 7.5, 300mM NaCl, 1mM TCEP, 10% (w/v) glycerol, and 1 mM ZnCl_2_. Human MLL2 (residues 5319–5537; accession # NM_003482), human MLL3 (residues 4689–4911; accession # NM_170606), and human MLL4 (residues 2490–2715; accession # NM_014727) were expressed in *E. coli* with N-terminal polyhistidine tags, and SET1B (residues 1629–1923; accession # NM_015048) was expressed in *E. coli* with N-terminal GST-tag. All four were assembled in complexes as MLL1, as mentioned above. Human recombinant NSD1 (residues 1538–2696; accession # NM_022455) was expressed with an N-terminal polyhistidine tag in an insect cell/baculovirus expression system. Human recombinant NSD2 (residues 2–1365; accession # NM_001042424) was expressed with an N-terminal polyhistidine tag in an insect cell/baculovirus expression system. Human recombinant NSD2-SET domain for SPR studies (residues 934–1241; accession # NM_001042424) was expressed with an N-terminal polyhistidine tag in Escherichia coli. Human recombinant NSD3 (residues 1021–1322; accession # NM_023034) was expressed in Escherichia coli as N-terminal GST fusions. Human PRDM9 (residues 2-414; accession # NM_020227) was expressed in an insect cell/baculovirus expression system as N-terminal GST fusions. Human recombinant PRMT1 (residues 2–371; accession # NM_001536) was expressed in Escherichia coli as N-terminal GST fusions. Human recombinant PRMT3 (residues 2–531; accession # NM_005788) was expressed in Escherichia coli as N-terminal polyhistidine tag. Human recombinant PRMT4 (residues 2–608; accession # NM_199141) was expressed in Escherichia coli as N-terminal GST fusion. Human recombinant PRMT5 (residues 2–637; accession # NM_006109) was expressed with an N-terminal Flag tag in an insect cell/baculovirus expression system. The complex of PRMT5/MEP50 was coexpressed PRMT5 above with MEP50 (residues 2–342; accession # NM_024102) with an N-terminal polyhistidine tag in an insect cell/baculovirus expression system. Human recombinant PRMT6 (residues 2–375; accession # NM_018137) and PRMT7 (residues 2–692; accession # NM_019023.2) were expressed as N-terminal polyhistidine tag in an insect cell/baculovirus expression system. Human recombinant PRMT8 (residues 61–394; accession # NM_019854) was expressed in Escherichia coli as N- and C-terminal polyhistidine tags. Human recombinant SET7/9 (residues 2–366; accession # NM_030648) was expressed in Escherichia coli as N-terminal GST fusion and C-terminal polyhistidine tag. Human recombinant SETD2 (residues 1434–1711; accession # NM_014159) and SMYD1 (residues 2–490; accession # NM_198274), both with N-terminal GST fusions, were expressed in *E. coli*. Human recombinant SMYD2 (residues 2–433; accession # NM_020197) was expressed in Escherichia coli as N-terminal polyhistidine tag. Human recombinant SUV39H1 (residues 44–412; accession # NM_003173) and SUV39H2 (residues 48–410; accession # NM_001193424), both with C-terminal polyhistidine tags, were expressed in *E. coli*. Human recombinant SUV420H1-tv2 (transcript variant 2, residues 2–393; accession # NM_016028) was expressed with an N-terminal GST fusion in an insect cell/baculovirus expression system. Purified nucleosomes (HMT-35-130) were obtained from HeLa according to Schnitzler (81). Core Histones, including histone 5 (HMT-35-435) purified from chicken erythrocytes by a modification of the method of Schnitzler (81) followed by acid extraction/dialysis (82). Human recombinant Histone H2A (residues 1–130; accession # NM_021052) and Histone H3.3 (residues 1-136; accession # NM_002107), both untagged, were expressed in *E. coli*. Human recombinant GST-GAR (Glycine and Arginine Rich sequence from the N-terminus of fibrillarin, residues 2–78; accession # NM_001436) was expressed in Escherichia coli as N-terminal GST fusion. The following substrates were purchased from vendors; Poly (dl-dC)(dl-dC) from Sigma-Aldrich, Lambda DNA from New England BioLabs, Histone H4 from BPS Biosciences, and Histone H3 (1-21) peptide from AnaSpec. The following reference compounds were purchased from vendors; SAH and Chaetocin from Cayman Chemicals, and LLY 507 and Ryuvidine from R & D Systems.

### Methyltransferase-Glo Assay

MTase-Glo assays were performed in a multi-step format in white, solid bottom 1,536-well plates (Greiner, Cat# 789175-F). First, 23 nL of compounds (or DMSO control) were pin-transferred into 3 μL of reaction buffer (50 mM Tris-HCl, pH 8.8, 5 mM MgCl_2_, 50 mM NaCl, 1 mM TCEP and 0.01% Tween 20) containing 666.7 nM [500nM final] nucleosomes and either 10.7 nM [8 nM final] wild-type, 16 nM [12 nM final] E1099K or 6.67 nM [5 nM final] T1150A NSD2 enzyme or no enzyme (low activity control, columns 2-3). Plates were then incubated for 30 min at room temperature prior to reaction initiation with 1 μL of 4 μM [1 μM final] S-adenosyl-L-methionine (SAM) in reaction buffer and incubated at room temperature for 15 min. Upon completion, methyltransferase conversion of SAM to SAH was then detected using a two-step detection system where: 1 μL of MTase-Glo Reagent was added to each well to convert SAH to ADP for 30 min at room temperature. Finally, 5 μL of MTase-Glo Detection Solution was added to each well and allowed to incubate for 30 min at room temperature to convert ADP to ATP, which was then measured by luminescence detection using a ViewLux uHTS Microplate Imager (PerkinElmer) and compared to control samples to determine relative activity.

### Methyltransferase-Glo Counterassay

MTase-Glo counterassay was performed with identical procedures as with the MTase-Glo primary assay but without NSD2 enzyme or nucleosomes. Instead 200 nM SAH was added to mimic the reaction with 20% substrate conversion.

### EPIgeneous HTRF Methyltransferase Assay

The EPIgeneous HTRF Methyltransferase assay was performed using, solid bottom white 1,536-well plates (Greiner, Cat# 789175-F). First, 23 nL of compounds (or DMSO control) were pin-transferred into 3 μL of reaction buffer (50 mM Tris-HCl, pH 8.8, 5 mM MgCl_2_, 50 mM NaCl, 1 mM TCEP and 0.01% Tween 20) containing 666.7 nM [500nM final] nucleosomes and either 10.7 nM [8 nM final] wild-type NSD2 enzyme or no enzyme (low activity control, columns 2-3). Plates were then incubated for 30 min at room temperature prior to reaction initiation with 1 μL of 4 μM [1 μM final] S-adenosyl-L-methionine (SAM) in reaction buffer and incubated at room temperature for 15 min. After the incubation period, 0.8 μL of EPIgeneous Detection Buffer One were added to each well, followed by a 10 min incubation at room temperature. Next, anti-SAH-Lumi4-Tb solution was prepared according to the manufacture’s instructions and 1.6 μL of the solution were added to each well. Finally, the SAH-d2 conjugate was prepared as a 32-fold dilution according to the manufacture’s instructions and 1.6 μL of the solution were added to each well. The assay plate was allowed to incubate for 1 hour at room temperature before detection of the HTRF signal using an Envision plate reader (PerkinElmer).

### Amplex Red (10-Acetyl-3,7-dihydroxyphenoxazine) Assay

The assay was adapted from a previously described protocol to assess redox cycling of compounds in the presence of reducing agents (48). 23 nL of compounds (or DMSO control) were pin-transferred into 2.5 μL HBSS (Thermo Fisher; containing 1.26 mM CaCl_2_, 0.49 mM MgCl_2_, 1 g/L D-glucose) in blacked walled 1,536 well plates. Compound fluorescence was measured immediately (READ 0) using a ViewLux uHTS microplate imager (PerkinElmer) equipped with Ex: 525/20 and Em: 598/25 filters. 2.5 μL of a 2X Amplex Red solution [100 μM Amplex Red (Cayman Chemical, Ann Arbor), 200 μM DTT (Thermo Fisher) and 2 U/mL horse radish peroxidase (Sigma-Aldrich); diluted in HBSS and protected from light] was added to each well. Fluorescence was measured after a 15 min incubation at room temperature (READ 1), using ViewLux settings identical to read 0. Activity was calculated using corrected fluorescence values (READ 1 minus READ 0), which were compared to control samples (negative = vehicle; positive = 46 μM walrycin B).

### HotSpot Methyltransferase Assay

The HotSpot radioisotope-based methyltransferase assays were performed as described previously (83, 84) with the following modifications. Standard substrate concentrations were 5 μM peptide or protein substrate, or 0.05 mg/ml for nucleosomes and core histones, and 1 μM SAM, unless otherwise mentioned. For control compound IC_50_ determinations, the test compounds were diluted in DMSO, and then added to the enzyme/substrate mixtures in nanoliter aliquots by using an acoustic technology (Echo550; Labcyte) with 20 min pre-incubation. The reaction was initiated by the addition of [^3^H]-SAM (tritiated SAM, PerkinElmer), and incubated at 30 °C for 1 hour. The reaction was detected by a filter-binding method. Data analysis was performed using GraphPad Prism software.

### Surface Plasmon Resonance

The Surface Plasmon Resonance measurement was performed using Biacore 8K (GE Healthcare) at 25 °C. Human recombinant NSD2-SET domain was immobilized to a Serial-S CM5 Sensorchip (GE Healthcare) using the classic amine-coupling method in the immobilization buffer containing 10 mM HEPES, 150 mM NaCl, 0.5 mM TCEP and 0.05% v/v Surfactant P20. Single cycle kinetic measurement were performed in the running buffer containing 50 mM Tris pH 8.8, 50 mM NaCl, 0.5 mM TCEP, 5 mM MgCl_2_, 0.05% Surfactant P20 and 2% DMSO. Compounds were diluted in a 3-fold manner in the running buffer and the DMSO concentration was carefully matched to 2%. Compound solutions were then injected over the prepared sensorchip at a flow rate of 80 μL/min for 80 seconds and allowed for dissociation period of 200-600 seconds. Data analysis was performed at Biacore 8K Evaluation software using the 1:1 kinetic binding model.

### Methyltransferase Profiling

The methyltransferase profiling was performed in HotSpot Methyltransferase assay format as described above. The final DMSO concentration in the reaction was adjusted at 1% DMSO for all profiling. The reaction buffer for EZH1 and EZH2 was 50 mM Tris–HCl, pH 8.0, 0.01% Brij35, 1 mM EDTA, 1 mM DTT, and 1 mM PMSF. The reaction buffer for SET8 and PRMT5 was 50 mM Tris–HCl, pH 8.5, 0.01% Brij35, and 1 mM DTT. The reaction buffer for NSD3 was 50 mM Bicine, pH 8.5, 0.01% Brij35, and 1 mM DTT. The reaction buffer for all other HMTs was 50 mM Tris–HCl, pH 8.5, 50 mM NaCl, 5 mM MgCl_2_, 1 mM DTT, and 1 mM PMSF. The reaction buffer for DNMTs was 50 mM Tris–HCl, pH 7.5, 5 mM EDTA, 0.01% Brij35, 5 mM DTT, 0.1 mM PMSF, and 5% glycerol.

### Western Blot Assay

Test compounds were dissolved with DMSO to a 10 mM stock. U-2 OS cells were seeded in 12-well plates at a density of 0.5 × 10^6^/well in complete culture medium and placed into the incubator at 37 °C, 5% CO_2_. After overnight incubation, the cells were treated with test compounds (10-dose with 3 fold dilution, 0.0025 - 50 μM), or with reference compound DZnep (10 μM single dose) and allowed to incubate for an additional 96 hours. Following the incubation with compound, culture media was removed and the cells were washed once with ice cold PBS. The cells were lysed with 1x SDS sample buffer (62.5 mM Tris-HCl pH 6.8, 2% SDS w/v, 10% Glycerol, 0.01% w/v bromophenol blue, 50 mM DTT) and the lysates were sonicated 3 times in 3 second increments at 15 amperes. Cell lysate samples (14 μL) were subjected to SDS-PAGE and transferred onto nitrocellulose membranes by the iBlot dry blotting system. The membranes were blocked with 2% non-fat milk blocking buffer for 1 hour and then probed with anti-histone H3 (Dimethyl Lys36) primary antibody overnight. Anti-rabbit IgG IRDye 680RD secondary antibody was used to detect the primary antibody. Then the blots were washed for 3 times with 1x TBS buffer plus 0.01% Tween 20 and re-probed with anti-histone H3 primary antibody and anti-mouse IgG IRDye 800CW secondary antibody. The membranes were scanned with a LI-COR Odyssey Fc Imaging System. The specific bands of interest were quantified by LI-COR Image Studio Lite software.

### U-2 OS Cell Growth Assay

U-2 OS (ATCC HTB-96) cells were obtained directly from ATCC and cultured according to the recommended culturing conditions. At cell passage 2, 1,750 cells were plated in 40 μl volumes (McCoy’s 5A + 10% FBS + 0.5x pen/strep) in a 384-well cell carrier plate (PerkinElmer) and incubated overnight. At 16 hours, test compounds were delivered in 195 nL aliquots by pin transfer. The plates were sealed with Breathe-Easy sealing membrane (Sigma) and images were captured every 4 hours up to 96 hours with an IncuCyte ZOOM System (Essen BioScience).

### Data Analysis

Data normalization and curve fitting were performed using in-house informatics tools. Briefly, raw plate reads for each titration point were first normalized relative to the DMSO-only wells (100% activity) and no enzyme control wells (0% activity), and then corrected by applying a plate-wise block pattern correction algorithm to remove any plate edge effects and systematic background noise. Active compounds from the primary HTS were defined as having a maximum response ≥ 50%. To determine compound activities from the 11-point qHTS, the concentration-response data for each sample was plotted and modeled by a four parameter logistic fit yielding IC_50_ and efficacy (maximal response) values as previously described (43). The activities were designated as class 1–4 according to the type of concentration-response curve observed. Active compounds were defined as having concentration-response curves in the classes of 1-3. The promiscuity score for each compound was defined as (number of assays that the compound is active)/(total number of assays that the compound was tested in). A compound with promiscuity score higher than 0.2 was considered as a “frequent hitter” to be eliminated from the follow-up studies.

## Acknowledgments

This work was supported by the NCATS Division of Pre-Clinical Innovation Intramural Program.

## Author Contributions

N.P.C., S.C.K., K.B., A.S., H.M., A.J. and M.D.H. conceived and initiated the research; N.P.C., S.C.K., M.J.H., O.W.L., K.Y.H., Y.W., Q.C., E.K., J.W., D.M.C., K.C-C.C, P.S., K.R.B., M.S., A.S., H.M., A.J., and M.D.H. conducted the research; all authors analyzed and discussed the data and contributed to and approved of the final manuscript.

